# Chronic alcohol intake elicits distinct multi-omic profiles in the liver *versus* skeletal muscle of mice

**DOI:** 10.1101/2025.10.09.681415

**Authors:** Craig R. G. Willis, Muni Swamy Ganjayi, Austin M. Brown, Samantha E. Moser, Nathaniel J. Szewczyk, Brian C. Clark, Cory W. Baumann

## Abstract

Alcohol-related liver disease and alcohol-related myopathy are widespread consequences of chronic alcohol use. However, understanding of the associated molecular mechanisms and effective treatments remains limited. To address this, we employed multi-omics to uncover molecular blueprints of liver *versus* skeletal muscle responses to chronic alcohol exposure, using a pre-clinical mouse model showing signs of alcohol-related liver dysregulation (diminished liver phosphatidylcholine-to-phosphatidylethanolamine lipid ratio) and alcohol-related myopathy (reduced muscle mass and strength). We found that the liver was more sensitive to chronic alcohol than muscle across the transcriptome, proteome and metabolome levels, but both tissues were equally sensitive at the lipidome level. The liver displayed an extensive and multi-layered metabolic molecular profile, while muscle was associated with upregulated inflammatory and matrisome responses and impaired mitochondrial energetics. Lipidome analyses also revealed a novel potential role for altered phospholipid remodelling in the aetiology of alcohol-related myopathy. Finally, computational drug repurposing identified several compounds for therapeutic targeting of alcohol-induced liver (e.g., saracatinib, GSK126) and muscle (e.g., metformin, trichostatin A) pathophysiology, perhaps working partly to counter metabolic dysregulation. Overall, our study provides a tractable list of therapeutic targets and treatments to help expedite the understanding of and countermeasures against alcohol-related liver disease and alcohol-related myopathy in humans.

## Introduction

Alcohol use remains widespread throughout society. In 2016, nearly a third of the world’s population were current alcohol drinkers (i.e., any alcoholic drinks in the past 12 months), with the average amount of alcohol consumed globally being ∼1.2 standard drinks daily^1^. By 2030, the global proportion of people drinking alcohol is expected to reach 50%^2^. Light-to-moderate alcohol intake has been linked to reduced risk of diabetes and cardiovascular disease^3,4^, however the ‘true’ health benefits of small quantities of alcohol remain contentious and are offset by an increased risk of other diseases (e.g., cancer) from modest alcohol consumption^5^. It is also well established that chronic, heavy alcohol intake is associated with many negative health outcomes. Over 200 health conditions, spanning multiple organ systems, are attributable to alcohol^6^. Excessive alcohol consumption is a leading contributor to the global burden of disease^7^ and a major risk factor for mortality^8^. In the US alone, excessive alcohol use contributes to approximately 13% of deaths among 20-to-64 year olds^9^ and carries an annual healthcare cost of $28 billion^10^. An ongoing priority of the World Health Organization is, therefore, to protect public health by preventing and reducing harmful use of alcohol^11^.

The liver is the main site for alcohol metabolism, making it one of the most afflicted organs by chronic alcohol intake^12^. Excessive alcohol use can lead to alcohol-related liver disease (ALD), which presents a wide clinical spectrum that includes alcohol-related fatty liver (steatosis), alcohol-related hepatitis, liver cirrhosis and liver cancer^13^. Alcohol-related fatty liver occurs in up to 90% of heavy alcohol drinkers and, while reversable with abstinence, can progress to more severe liver disease (e.g., cirrhosis) and increase the risk of liver-related mortality^14–16^. Liver cirrhosis is the 11^th^ leading cause of death worldwide and develops in 10-20% of heavy alcohol users, with 30-50% of global cirrhosis-related deaths attributable to alcohol^17,18^. However, despite ALD being the most prevalent type of chronic liver disease globally^19^ and the leading cause of alcohol-specific death in some countries (e.g., the UK^20^), therapeutic options are still lacking. Abstinence can be challenging for alcohol-dependent individuals and cannot reverse advanced stages of ALD^21,22^. Nevertheless, currently, there no approved pharmacologic treatments for ALD^18,23,24^.

Another major organ impacted by alcohol is skeletal muscle. Excessive alcohol intake can cause muscle atrophy and weakness, a condition known as alcohol-related myopathy^25^. Skeletal muscle is the largest tissue in the human body^26^ and is central to many key physiological processes, including locomotion and metabolism^27^. Alcohol-induced muscle atrophy can therefore have profound negative implications on health and physical function, ultimately reducing quality of life^28^. Strength loss can exceed 30% in chronic, heavy alcohol drinkers and is only partially restored with abstinence^29^. With alcohol-related myopathy affecting 45-70% of heavy drinkers, it is one of the most widespread alcohol-related conditions^30^. Nevertheless, the exact pathophysiologic mechanisms of alcohol-related myopathy remain incompletely defined and likely multifactorial, with impaired muscle protein synthesis, increased muscle protein degradation, mitochondrial dysfunction, perturbed metabolism, inflammation, oxidative stress and diminished muscle regenerative capacity all purported to contribute^28^. Moreover, like ALD, therapeutic strategies for alcohol-related myopathy are limited^31^.

The development and progression of ALD and alcohol-related myopathy are likely caused not only by the direct effects of alcohol on liver (ALD) and muscle (alcohol-related myopathy), but also via indirect effects across these two tissues. Indeed, ALD may contribute to alcohol-related myopathy, and *vice versa*, by influencing signalling between the liver and skeletal muscle^28,30,32–34^. Certain alcohol-induced phenotypes can also be common between the liver and muscle, such as tissue fibrosis^13,35^. Thus, ALD and alcohol-related myopathy encompass distinct, shared, *and* interlinked pathophysiologic hallmarks. Understanding similarities and differences in molecular responses of the liver and skeletal muscle to chronic alcohol exposure could, therefore, expedite the development of more targeted therapeutic interventions for both conditions. However, the molecular dynamics of excessive alcohol consumption in the context of liver *versus* skeletal muscle remain largely unknown.

Molecular and translational medicine have been revolutionised by the paradigm shift of biology into a ‘big data’ era. Indeed, modern ‘omics’ technologies now allow for the characterisation of the molecular milieu at an immense resolution^36^, paving the way for unprecedented medical advances from identifying individuals with undiagnosed disease to guiding personalised medicine^37^. The omics cascade includes several layers that bridge genotype to phenotype, from genomics and transcriptomics through to proteomics and metabolomics/lipidomics^38^. Research efforts have often focussed on just a single omics layer at a time, overlooking the fact that complex disease pathophysiology is underpinned by disturbances across multiple layers of molecular biology and their interplay^39^. The integration and interrogation of different omics data types, termed ‘multi-omics’, can improve biological insights by providing a more holistic ‘systems-level’ view of disease aetiology^40^. In turn, multi-omics offers an unparalleled platform for aiding the discovery and development of new disease diagnostics, prognostics, treatments, and preventative strategies^41^.

Here, we harnessed the power of multi-omics to define and compare the molecular blueprints of chronic alcohol use in the liver and skeletal muscle using an established pre-clinical mouse model. Our findings provide valuable insights into the molecular mechanisms underlying alcohol-induced liver and skeletal muscle pathophysiology, and reveal new therapeutic targets and candidate drug options. Consequently, the results from this innovative multi-omics study provide a strong foundation for accelerating the understanding of, and targeted treatments against, ALD and alcohol-related myopathy in humans.

## Results

### Mice that drink alcohol chronically show signs of alcohol-related myopathy and ALD

Transcriptomic, proteomic, metabolomic and lipidomic data were generated from the liver and plantarflexor muscles (*gastrocnemius*, *plantaris*, *soleus*) of adult C57BL/6 female mice that had consumed either 100% water (Control mice, *n* = 9) or 80% water + 20% alcohol (ethanol) (Alcohol mice, *n* = 14) for 34-40 weeks^42,43^. Mice that consumed alcohol for 34-40 weeks had elevated blood alcohol levels (Control = 0 (3.1) mg.dL^-1^, Alcohol = 159.3 (120.6) mg.dL^-1^; *P* = 7.60E-05) and lower total body mass (Control = 28.1 (6.5) g, Alcohol = 25.8 (3.7) g; *P* = 5.55E-03), fat mass (Control = 7.0 (2.1) g, Alcohol = 5.4 (1.9) g; *P* = 1.73E-02) and lean mass (Control = 18.5 (2.5) g, Alcohol = 17.0 (1.1) g; *P* = 1.34E-03) compared to controls. Plantarflexor muscle mass (Figure 1A) and *in vivo* isometric torque (Figure 1B) were lower in alcohol-consuming mice relative to control mice, consistent with alcohol-induced muscle atrophy and weakness (i.e., alcohol-related myopathy). Alcohol-consuming mice also had an elevated ratio of phosphatidylcholine (PC) to phosphatidylethanolamine (PE) lipid content in muscle (marker of muscle insulin resistance^44^) (Figure 1C), demonstrating that muscle metabolism may have changed after alcohol intake^25^. While liver mass was similar between control mice and alcohol-consuming mice (Figure 1D), chronic alcohol drinking lowered hepatic PC:PE ratio values (Figure 1E), which is indicative of diminished hepatic membrane potential and a strong indicator of ALD in both pre-clinical models and human patients.^45^

**Figure 1:**
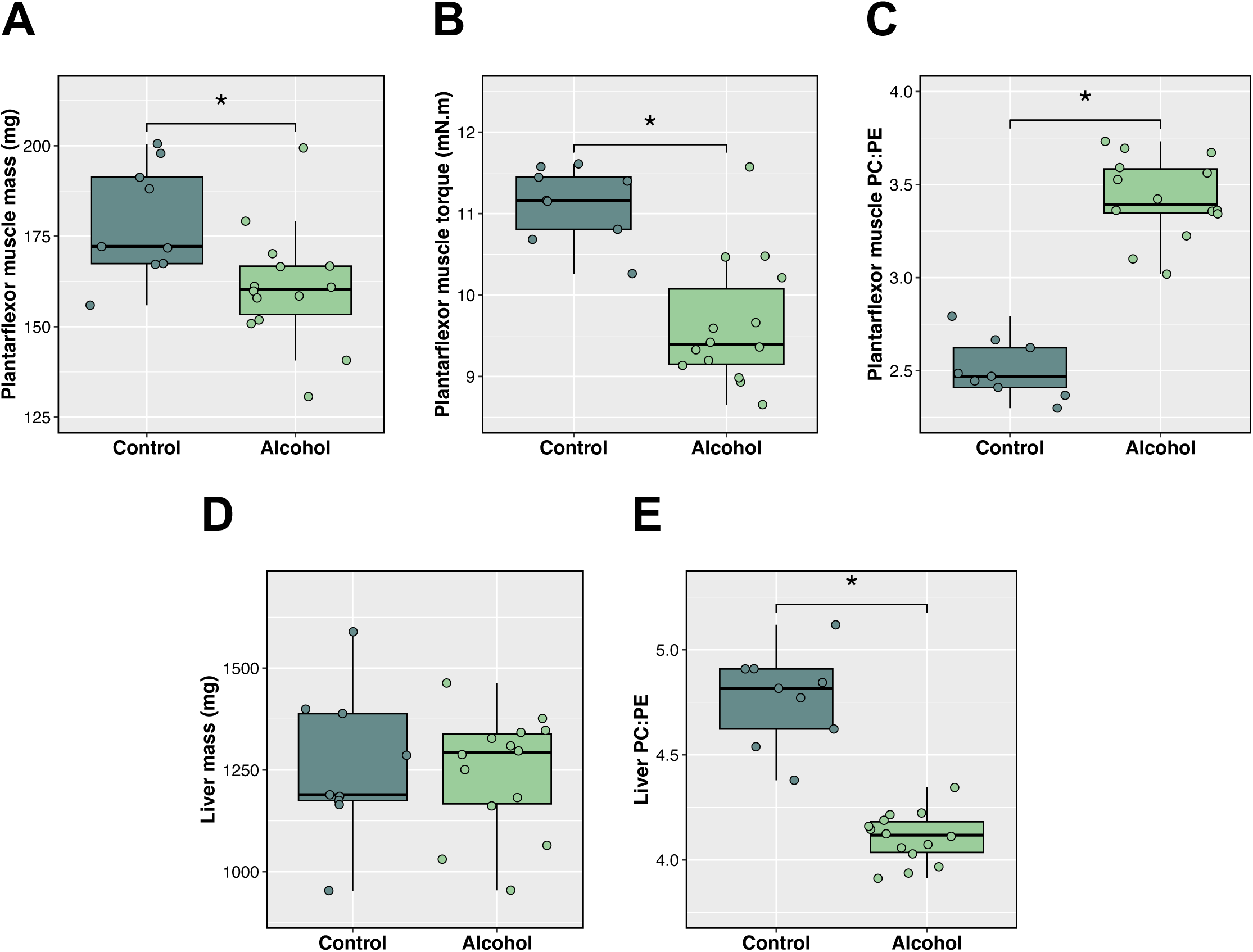
Indicators of alcohol-related myopathy and ALD following chronic alcohol consumption. Boxplots illustrate plantarflexor muscle mass (Panel **A**), plantarflexor muscle peak isometric contractile torque (Panel **B**), plantarflexor muscle PC:PE ratio (Panel **C**), liver mass (Panel **D**) and liver PC:PE ratio (Panel **E**) in alcohol-consuming mice *versus* control mice. Analyses via two-tailed Student’s/Welch’s independent *t*-test or Mann-Whitney *U* test, as appropriate. *: *P* ≤ 0.05.

### Chronic alcohol consumption extensively disrupts liver and muscle transcriptomes

Chronic alcohol markedly perturbed the transcriptomes of both liver and muscle, with over 1,000 dysregulated genes in each case (Figure 2A). However, the liver transcriptome was more sensitive to the effects of chronic alcohol, exhibiting more differentially expressed genes and greater magnitudes of change compared to skeletal muscle (Figure 2A). An overlay of dysregulated genes showed that liver and muscle experience largely unique transcriptomic responses to chronic alcohol (Figure 2B). In the liver, uniquely upregulated genes were enriched with MTA3 transcription factor (TF) targets and glycolysis genes, while uniquely downregulated genes were associated with processes such as cholesterol homeostasis and hypoxia (Figure 2C, Figure S1). In muscle, uniquely upregulated genes largely mapped to inflammatory processes, apoptosis and extracellular matrix processes as well as CRELD1 and LYL1 TF targets, while uniquely downregulated genes were linked to fatty acid metabolism, oxidative phosphorylation and mitochondrial biogenesis, and included targets of the NR1H2 and PPARGC1 TFs (Figure 2C, Figure S1). Fewer genes were commonly dysregulated by chronic alcohol across both tissues, with commonly upregulated gene involved in carbohydrate metabolism processes, and commonly downregulated genes related to amino acid metabolism, chromosome-related processes, and targets of the AFF4 and RXRG TFs (Figure 2C, Figure S1). Tissue specificity was also evident among the top genes dysregulated by alcohol, with virtually all top genes in liver (except *Arsa*, *Sox12* and *Paqr7)* unaffected by alcohol in muscle and, similarly, nearly all top genes in muscle (except *Gstk1*, *Ivd*, *Pxmp2* and *Sertad3*) unaffected by alcohol in the liver (Figure 2A). Top liver-specific genes included many related to alcohol and/or cholesterol metabolism (*Cyp2d9*, *Cyp3a41b*, *Cyp3a44*, *Cyp51*, *Idi1*, *Msmo1*, *Nsdhl*, *Rdh11*, *Tkfc*), a glutathione conjugator (*Gstm3*; upregulated) and a leptin receptor (*Lepr*; upregulated). Top muscle-specific genes included those linked to cytoskeletal regulation (*Actc1*, *Tmsb10*, *Vasp*; all upregulated), metabolism of pro-inflammatory mediators (*Cyp4f18*; upregulated) and mitochondrial energetics (*Coq10a*, *Mpc1*; both downregulated) (Figure 2A).

**Figure 2:**
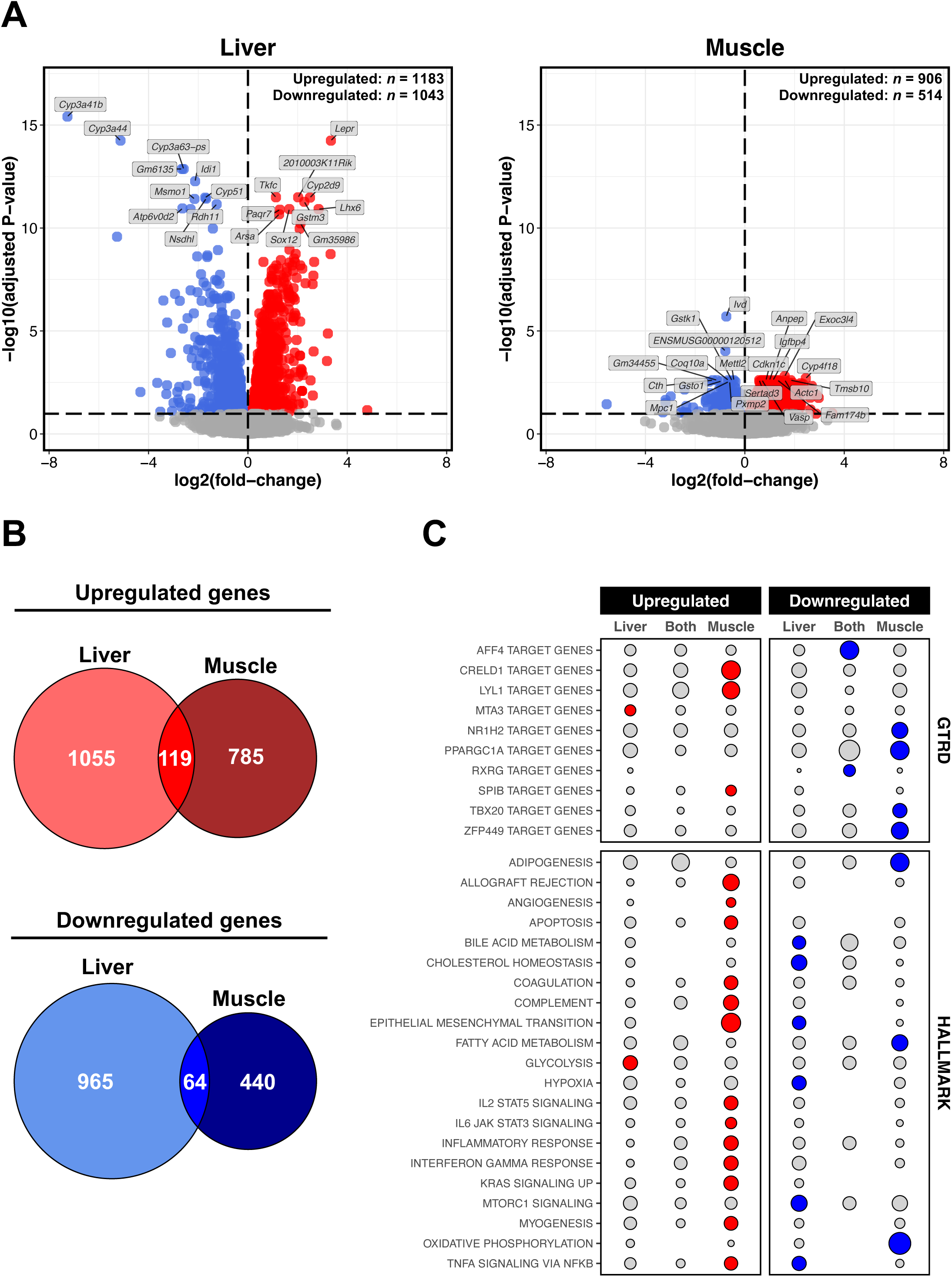
Liver *versus* muscle transcriptomic responses to chronic alcohol. Panel **A**: Volcano plots for differential gene expression analysis (Alcohol *versus* Control) in the liver and muscle. Red and blue shading denote significant upregulation and downregulation, respectively (adjusted *P* ≤ 0.1). Annotated genes are those ranked in top 10 upregulated/downregulated based on *t*-score. Panel **B**: Venn diagrams illustrating degree of overlap between genes upregulated/downregulated by chronic alcohol use in the liver and muscle. Panel **C**: Bubble plot depicting results from over-representation analysis of MSigDB mouse Molecular Hallmark and GTRD gene sets for genes commonly/uniquely dysregulated by chronic alcohol use in the liver and muscle (as per panel **B**). Circle size is proportional to the number of gene hits as a % of the total number of annotated genes for a given overlap. Red and blue shading denote significant over-representation in upregulated and downregulated genes, respectively (adjusted *P* ≤ 0.05, enriched for ≥ 2 genes).

### Dysregulation of the muscle proteome is minimal following chronic alcohol intake

Like the transcriptome, the liver proteome was more sensitive to chronic alcohol than the muscle proteome (Figure 3A). While approximately 600 proteins were dysregulated by alcohol in liver, only 8 proteins were dysregulated in muscle (Figure 3A). Overlaying these signatures revealed a major liver-specific proteome response to chronic alcohol, accompanied by a limited muscle-specific response and few commonly dysregulated proteins (Figure 3B). In liver, uniquely upregulated proteins were involved in processes such as oxidative phosphorylation, lipid metabolism, and mitochondrial translation, while uniquely downregulated proteins were associated with cholesterol homeostasis, glycolysis and amino acid metabolism (Figure 3C). In contrast, proteins uniquely upregulated in muscle were enriched in processes such as coagulation, haemostasis, and the immune complement system (including the direct terminal complement pathway inhibitor, clusterin^46^, and the classical and alternative complement pathway modulator, antithrombonin-III^47^) (Figure 3C). No proteins were uniquely downregulated in muscle (Figure 3B), neither were there any enriched terms for commonly dysregulated proteins nor enriched TF target sets in any case (Figure 3C). Unsurprisingly, all top-dysregulated proteins in the liver were unaffected by alcohol in muscle (Figure 3A), with most (except Alg3, Fcn1, Galt, Gnaq, Med,16, Psmb9) also being uniquely dysregulated at the gene level in the liver. This included several metabolism-related proteins (Cbr3, Dcxr, Idi1, Lpcat3, Mvk, Pfkfb1, Retsat) and two glutathione conjugators (Gstm1, Gstm3; both upregulated) (Figure 3A). In contrast, top muscle-specific proteins included stress-related chaperones (Bcap29, Clu; both upregulated), blood coagulation regulators (Serpinc1; upregulated) and immune/inflammatory elements (Cd5l, Mbl1; both upregulated) (Figure 3A).

**Figure 3:**
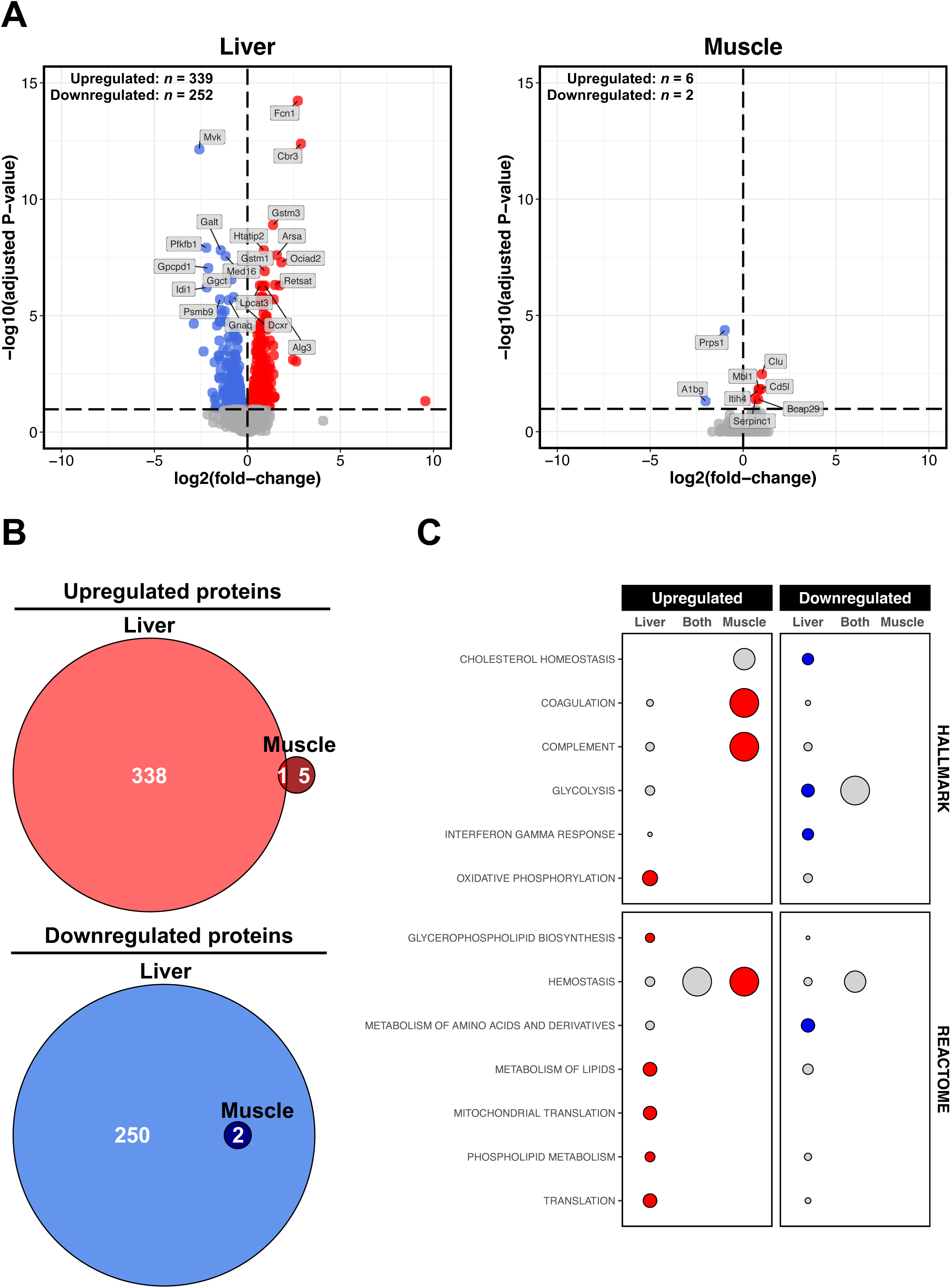
Liver *versus* muscle proteomic responses to chronic alcohol. Panel **A**: Volcano plots for differential protein abundance analysis (Alcohol *versus* Control) in the liver and muscle. Red and blue shading denote significant upregulation and downregulation, respectively (adjusted *P* ≤ 0.1). Annotated proteins are those significant proteins ranked in top 10 upregulated/downregulated based on *t*-score. Panel **B**: Venn diagrams showing degree of overlap between proteins upregulated/downregulated by chronic alcohol use in the liver and muscle. Panel **C**: Bubble plot depicting results from over-representation analysis of MSigDB Molecular Hallmark and Reactome Pathways gene sets for proteins commonly/uniquely dysregulated by chronic alcohol use in the liver and muscle (as per panel **B**). Circle size is proportional to the number of hits as a % of the total number of annotated genes for a given overlap. Red and blue shading denote significant over-representation in upregulated and downregulated proteins, respectively (adjusted *P* ≤ 0.05, enriched for ≥ 2 features).

### Transcriptome and proteome responses to chronic alcohol use are more synergistic in the liver compared to muscle

With our transcriptomic and proteomic data exhibiting some consistent outcome themes (e.g., quantitative differential patterns and functions of dysregulated features), we further explored the general concordance between gene and protein responses to chronic alcohol in each tissue using ‘threshold-free’ approaches to maximise global biological signal^48^. We observed a strong degree of agreement in the liver between the differential patterns of features present at both the gene and protein levels (*n* = 3,303 genes/proteins) (Figure 4A). While a trend for agreement between differential gene and protein patterns was also observed in muscle, the corresponding signal was much weaker compared to that in the liver (Figure 4A). Similarly, the agreement for global pathway regulation between gene and protein levels was more apparent in the liver than in muscle (Figure 4B). These analyses revealed numerous themes in line with our individual transcriptome and proteome analyses. For example, in muscle, there was a strong upregulation of inflammatory (e.g., ‘neutrophil degranulation’, ‘innate immune system’, ‘platelet aggregation signalling and aggregation’, ‘complement cascade’), extracellular matrix and apoptosis pathways. In the liver, there was a unique downregulation of cholesterol homeostasis pathways. Both liver and muscle showed common downregulation of amino acid metabolism pathways at the gene level, which extended only to the protein level in the liver. Additionally, there were opposing mitochondrial and oxidative phosphorylation responses in the liver compared to the muscle (Figure 4B).

**Figure 4:**
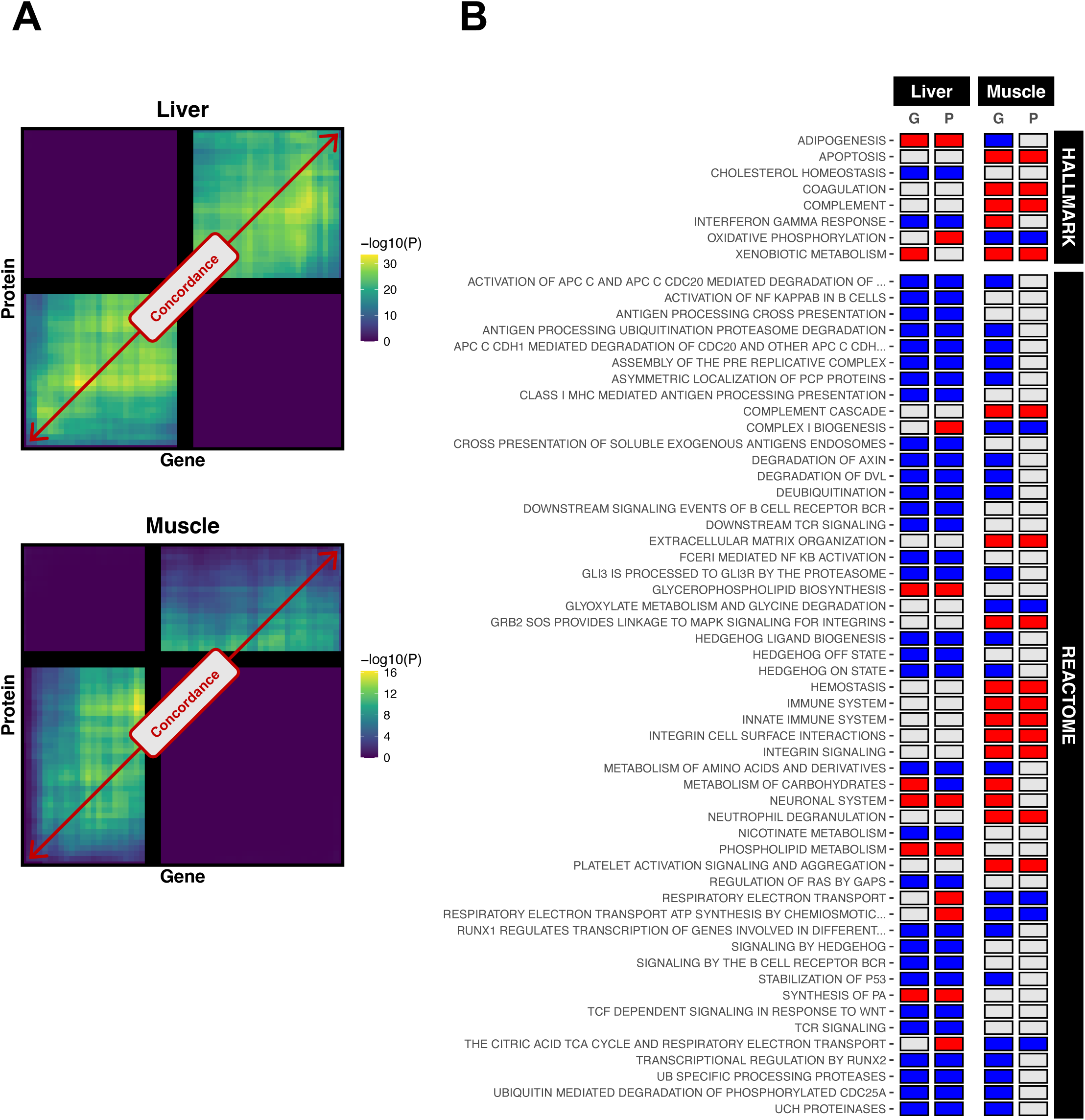
Global comparison between transcriptomic and proteomic responses to chronic alcohol in the liver and muscle. Panel **A**: Rank-rank hypergeometric overlap (RRHO) plots illustrating the degree of correspondence between gene level and protein level responses to chronic alcohol use in the liver and in muscle. The lighter the colouring in the lower-left quadrant, the stronger the concordance in upregulation between genes and proteins. The lighter the colouring in the upper-right quadrant, the stronger the concordance in downregulation between genes and proteins. For RRHO analysis, only unique features present at both the gene level and protein level were included (*n* = 3303 gene/proteins), with features ranked on *t*-score. Panel **B**: Heatmap comparing global pathway regulation by chronic alcohol use in liver and muscle at gene and protein levels. Shown are those MSigDB mouse Molecular Hallmarks and Reactome Pathways that are similarly regulated at gene and protein levels in at least one of the two tissues. Results were derived via gene set enrichment analysis applied to unique features present at both the gene level and protein level (*n* = 3303 gene/proteins), with features ranked on *t*-score. Red and blue shading denote significant upregulation (adjusted *P* ≤ 0.1, normalised enrichment score > 0) and downregulation (adjusted *P* ≤ 0.1, normalised enrichment score < 0), respectively. G = gene, P = protein.

### Metabolome responses to chronic alcohol are less pronounced in muscle than the liver

As at the transcriptome and proteome levels, the liver metabolome was more sensitive to chronic alcohol than the muscle metabolome, with six-fold more metabolites dysregulated in the liver compared to muscle (Figure 5A). Overlaying differential metabolites revealed a strong liver-specific response to alcohol, with fewer metabolites either uniquely dysregulated in muscle or commonly dysregulated both tissues (Figure 5B). In the liver, uniquely upregulated metabolites were strongly associated with the glycine, serine and threonine metabolism pathway, while uniquely downregulated metabolites were linked to the carbohydrate metabolism pathway and the organic acid class (Figure 5C). Metabolites downregulated by alcohol in both tissues were predominantly organic nitrogen compounds and metabolites involved in glycerophospholipid metabolism (Figure 5C). However, no enriched terms were identified for metabolites either upregulated in both tissues or uniquely dysregulated in muscle (Figure 5C). Interestingly, despite the general tissue specificity of upregulated metabolites (Figure 5B), liver and muscle shared many top-upregulated metabolites, including 2-deoxy-d-glucose, 3-o-beta-d-galactosyl-sn-glycerol, azithromycin, fructose (all carbohydrate related) and quinolinic acid (pyridine related) (Figure 5A). Acetylcholine was the only top-downregulated metabolite in both tissues, with all other top-downregulated metabolites in the liver being metabolites unaffected by alcohol in muscle (Figure 5A). Conversely, only 4 top-downregulated metabolites in muscle were not dysregulated in the liver, namely hecogenin (a terpenoid), cholfenethol (an acaricide), 4-tert-octylphenol monoethoxylate (a fatty alcohol) and beta-asarone (a phenylpropanoid with anti-inflammatory and anti-apoptotic potential^49^) (Figure 5A). Top liver-specific metabolites included the upregulated compound (9cis)-retinal (a retinol-related compound), and downregulated compounds such as propionylcarnitine (a fatty acid ester), bufotalin (a steroid), trans-3-indoleacrylic acid (an indole), hydroxyphenyllactic acid (a phenylpropanoic acid), beta-alanine and guanidinosuccinic acid (both amino acid-related compounds) (Figure 5A). The identification confidence levels for all analysed metabolites are detailed in Document S1 of the supplementary information. These range from Level 1 (highest confidence), assigned to metabolites definitively identified by matching authentic standards, to Level 4 (low confidence), representing unknown compounds or tentative assignments based on database matches without confirmatory spectral or standard-based validation.

**Figure 5:**
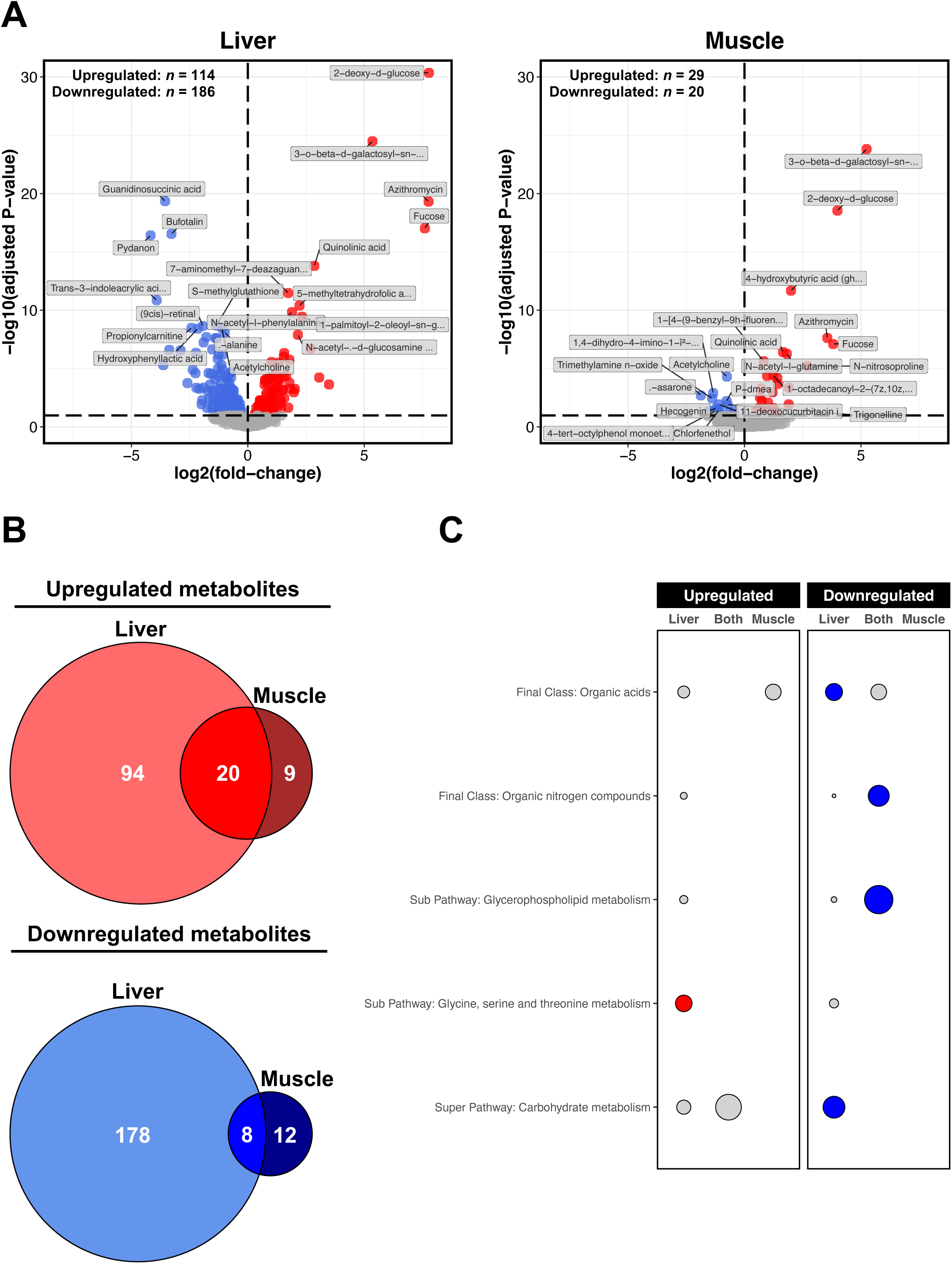
Liver *versus* muscle metabolomic responses to chronic alcohol. Panel **A**: Volcano plots for differential metabolite analysis (Alcohol *versus* Control) in the liver and muscle. Red and blue shading denote significant upregulation and downregulation, respectively (adjusted *P* ≤ 0.1). Annotated metabolites are those significant metabolites ranked in top 10 upregulated/downregulated based on *t*-score. Panel **B**: Venn diagrams showing degree of overlap between metabolites upregulated/downregulated by chronic alcohol use in the liver and muscle. Panel **C**: Bubble plot depicting results from over-representation analysis of metabolite final class, super pathway and sub pathway annotations for metabolites commonly/uniquely dysregulated by chronic alcohol use in the liver and muscle (as per panel **B**). Circle size is proportional to the number of metabolite hits as a % of the total number of annotated metabolites for a given overlap. Red and blue shading denote significant over-representation in upregulated and downregulated metabolites, respectively (adjusted *P* ≤ 0.05, enriched for ≥ 2 metabolites).

### Liver and muscle lipidomes are equally sensitive to chronic alcohol consumption

Unlike all other omics layers studied, the lipidomes of liver and muscle were comparably sensitive to chronic alcohol, with approximately 320 lipids differentially regulated in each case (Figure 6A). Nevertheless, overlaying lipidome profiles revealed large tissue-specific lipid responses to alcohol, although noticeable proportions of commonly dysregulated lipids were also found (Figure 6B). Commonly upregulated lipids were enriched with P-ethanol compounds, while commonly downregulated lipids were enriched with P-choline compounds (Figure 6C). Lipids uniquely upregulated in the liver predominantly belonged to the P-inositol class, whereas those uniquely downregulated in the liver were enriched with sphingolipids (Figure 6C). Lipids uniquely upregulated in muscle strongly mapped to the P-ethanol Amine class, although no enriched classes were identified for lipids uniquely downregulated in muscle (Figure 6C). Among the top-downregulated lipids in muscle were P-choline, P-ethanol and P-ethanol Amine compounds that were either upregulated (Pc(36:6)(rep), Pet(16:0/20:3)) or unperturbed (Pc(36:6), Pc(44:12), Dmepe(40:6p)) by alcohol in liver (Figure 6A). Similarly, half of the top-upregulated lipids in muscle were compounds downregulated by alcohol in the liver (Pc(16:0/18:2), Pc(17:0/18:2), Pc(33:2), Pc(36:3), Ps(39:3)) (Figure 6A). In both liver and muscle, the top-upregulated lipid was Pet(16:0/18:2), a P-ethanol compound and potential alcohol biomarker in blood^50^ (Figure 6A).

**Figure 6:**
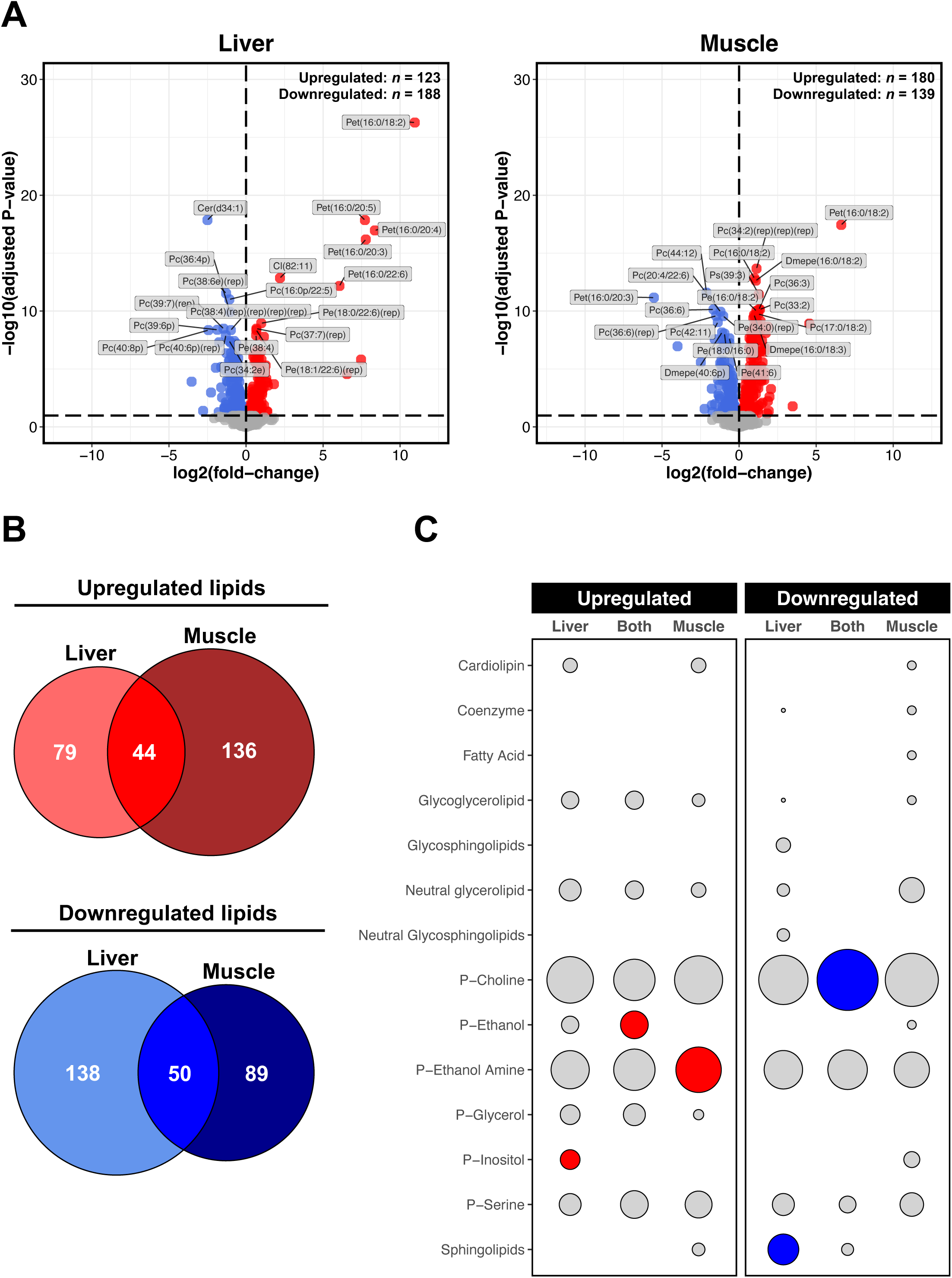
Liver *versus* muscle lipidomic responses to chronic alcohol. Panel **A**: Volcano plots for differential lipid analysis (Alcohol *versus* Control) in the liver and muscle. Red and blue shading denote significant upregulation and downregulation, respectively (adjusted *P* ≤ 0.1). Annotated lipids are those significant lipids ranked in top 10 upregulated/downregulated based on *t*-score. Panel **B**: Venn diagrams illustrating degree of overlap between lipids upregulated/downregulated by chronic alcohol use in the liver and muscle. Panel **C**: Bubble plot depicting results from over-representation analysis of main lipid classes for lipids commonly/uniquely dysregulated by chronic alcohol use in the liver and muscle (as per panel **B**). Circle size is proportional to the number of lipid hits as a % of the total number of annotated lipids for a given overlap. Red and blue shading denote significant over-representation in upregulated and downregulated lipids, respectively (adjusted *P* ≤ 0.05, enriched for ≥ 2 lipids).

### Multi-omic networks discriminate chronic alcohol use in the liver and in muscle

We also performed discriminative multi-omic network analysis to model complex relationships across the different omics layers of molecular biology^51^ and, in turn, establish further key molecular drivers of alcohol-induced muscle/liver pathophysiology^52^. In the liver, a multi-omic relevance network of 309 features was deduced, comprising a mix of genes (*n* = 187), proteins (*n* = 48), metabolites (*n* = 19) and lipids (*n* = 55). The liver relevance network partitioned into 4 sub-networks of varying size (Liver C1-C4), each containing at least one feature from each omics type (Figure 7A). Virtually all (*n* = 301) features of the liver relevance network showed dysregulation by chronic alcohol in the liver (Figure 7A). Hub analyses further defined a list of 40 ‘priority’ features across the liver network, including 11 genes, 8 proteins, 6 metabolites, 15 lipids (Figure 7A). Many of these upregulated features were related to metabolism and energy homeostasis, including: the genes *Ephx1* (lipid metabolism), *Pfkm* (glycolysis) and *Zfp385*a (adipogenesis) (Liver C2 hubs); the proteins Ak2 (energy homeostasis), Hadha (mitochondrial beta-oxidation), Htatip2 (redox sensor), Stbd1 (cargo receptor for glycogen) (Liver C1 hubs) and Pc (glucose and lipid synthesis) (Liver C4 hub), and; the metabolites 3-o-beta-d-galactosyl-sn-glycerol (carbohydrate metabolism), fucose (carbohydrate class) (Liver C1 hubs), 2-deoxy-d-glucose (carbohydrate class) and azithromycin (carbohydrate class) (Liver C4 hubs) (Figure 7A). Several upregulated features related to conjugation were also identified among liver hubs, including *Ugt1a9* (glucuronidation pathway) (Liver C2 hub) and the reduced glutathione conjugators *Gstp1* (Liver C2 hub) and Gstm1 (Liver C1 hub) (Figure 8A). Additionally, more than half of liver hub lipids were downregulated P-choline compounds, including Lpc(15:1), Pc(31:0), Pc(32:1e), Pc(39:7)(rep) (Liver C1 hubs), Pc(16:0p/22:5), Pc(38:4e), Pc(38:6e)(rep) (Liver C4 hubs), and the sole hub of Liver C3, Pc(36:4p) (Figure 7A). Pet(16:0/18:2) (P-ethanol compound), the top-upregulated lipid in liver (Figure 7A), was also found among the hubs of Liver C4 (Figure 7A).

**Figure 7:**
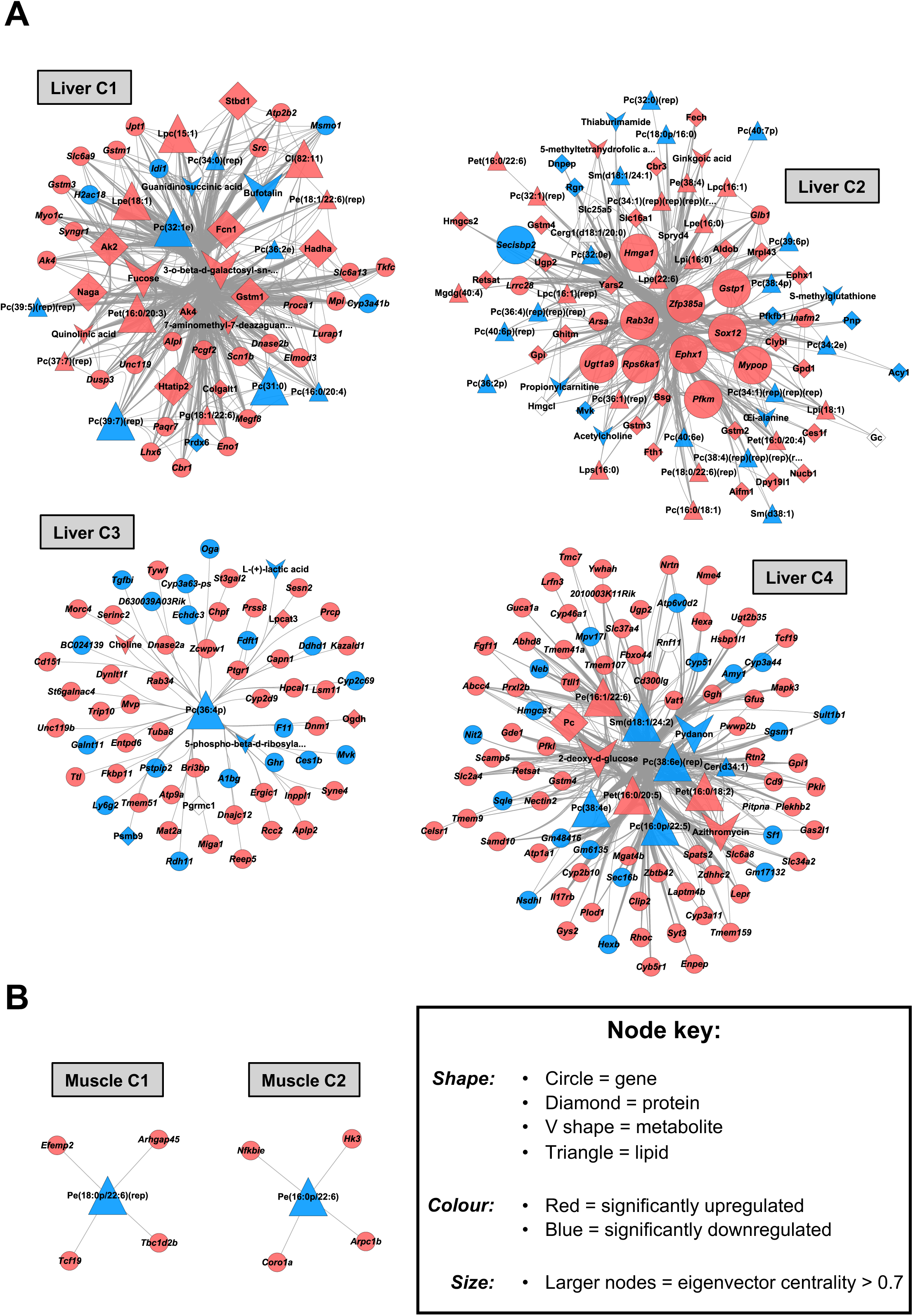
Integrative multi-omic network modelling of the liver and muscle responses to chronic alcohol. Communities of strongly connected molecular features in the liver (panel **A**) and muscle (panel **B**) with chronic alcohol use, as derived via Mixomics DIABLO analyses with downstream correlation network generation (|correlation coefficient| > 0.85, with component 1 used for the liver and component 2 used for muscle).

**Figure 8:**
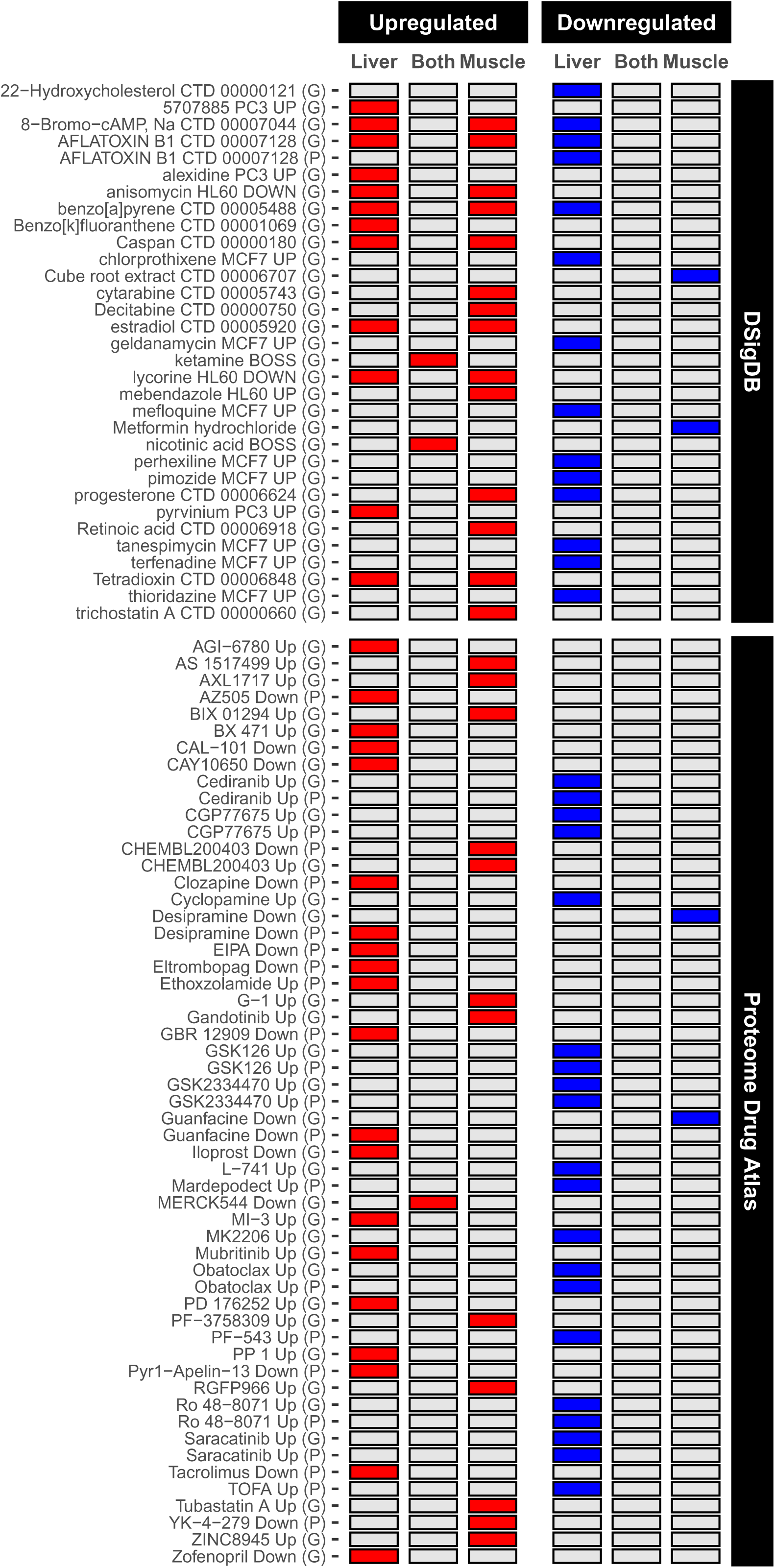
Omics-driven repurposing of drug therapeutics for ALD and alcohol-related myopathy. Heatmap includes results from over-representation analysis of human Drug Signature Database (DSigDB) and Proteome Drug Atlas sets for genes/proteins commonly/uniquely dysregulated by chronic alcohol use in the liver and muscle. Shown are the top 10 most significant sets for each common/unique feature permutation. Red and blue shading denote significant over-representation in upregulated and downregulated genes/proteins, respectively (adjusted *P* ≤ 0.05, enriched for ≥ 2 features). G = enriched when genes used as input, P = enriched when proteins used as input.

Compared to the liver, the muscle multi-omic relevance network was much smaller and contained only genes (*n* = 8; all downregulated by alcohol in muscle) and lipids (*n* = 2; both upregulated by alcohol in muscle). The muscle relevance network partitioned into 2 sub-networks (Muscle C1 and C2), each comprising 1 lipid and 4 genes (Figure 7B). Muscle C1 contained upregulated genes related to transcription (*Tcf19*), endocytosis (*Tbc1d2b*), collagen fibril assembly (*Efemp2*) and actin cytoskeleton regulation (*Arhgap45*), all centred around Pe(18:0p/22:6)(rep), a downregulated P-ethanol Amide compound and the sole hub feature within Muscle C1 (Figure 7B). Muscle C2 followed a similar topology, with upregulated genes related to NF-Kappa-B inhibition (*Nfkbie*), hexose phosphorylation (*Hk3*) and actin cytoskeleton dynamics (*Arpc1b*, *Coro1a*) all converging on one downregulated ‘hub’ P-ethanol Amide compound, Pe(16:0p/22:6) (Figure 7B).

### Chronic alcohol use associates with omic profiles that are targetable by drugs

Given the need to establish new therapeutics against ALD and alcohol-related myopathy, we screened for potential drug treatments in humans based on the omic signatures derived in this study. We specifically focused on transcriptomic and proteomic profiles due to the greater availability of drug-target annotations compared to the metabolome and lipidome. Consequently, a total of 745 compounds from the Drug Signature Database^53^ and Proteome Drug Atlas^54^ were predicted to target features dysregulated by chronic alcohol in liver and/or muscle. Consistent with the general tissue-specificity observed in transcriptome and proteome responses to alcohol, virtually all (*n* = 742) compounds were predicted to target features uniquely dysregulated by chronic alcohol in either liver or muscle. This trend remained evident even when considering the top drug compound predictions based on significance (Figure 8). Among the predicted compounds for liver-specific features, several were reproducibly identified across transcriptome and proteome inputs. These included Saracatinib and GSK126, which may potentially hold anti-liver fibrotic properties^55,56^ (Figure 8). Predicted compounds for features uniquely dysregulated by alcohol in muscle included metformin, trichostatin A and retinoic acid, each of which may positively impact the phenotype observed with alcohol-related myopathy^57–59^ (Figure 8). Only three compounds were predicted for features commonly upregulated by chronic alcohol in both liver and muscle, namely ketamine, nicotinic acid and MERCK544, while no compounds were predicted for features commonly downregulated by chronic alcohol in both tissues (Figure 8). However, several compounds, such as estradiol and lycorine, were predicted to target unique responses in both tissues (Figure 8), indicating that some compounds may impact both liver and muscle after alcohol intake, but through distinct mechanisms. The presence of carcinogenic factors among predicted compounds (e.g., AFLATOXIN B1, benzo[a]pyrene) (Figure 8) further underscores the toxic effects of chronic alcohol on both liver and skeletal muscle tissues.

## Discussion

Alcohol use is a leading cause of morbimortality worldwide^7,8^, such that minimising its harmful effects is a major public health priority of the World Health Organization^11^. Among the most serious and prevalent consequences of excessive alcohol consumption are ALD and alcohol-related myopathy^17,30^. However, therapeutic options are currently lacking, underscoring the urgency for continued research into the molecular blueprints of alcohol-induced liver and muscle pathophysiology. Multi-omics has emerged as a cutting-edge frontier for accelerating molecular understanding of and developing countermeasures against complex diseases^40,41^. Therefore, we undertook the first multi-omic screen of chronic alcohol signatures in liver *versus* skeletal muscle. Our findings revealed that liver and muscle are characterised by largely unique molecular profiles in the context of chronic, excessive alcohol consumption. Consequently, liver and muscle from mice that consumed alcohol were associated with mostly divergent therapeutic targets and candidate pharmacologic interventions. These results have important clinical implications for developing optimal strategies to ameliorate ALD and alcohol-related myopathy both individually and synergistically.

Mechanistic investigations of chronic alcohol drinking in people come with significant ethical and technical challenges^60^, highlighting the importance of pre-clinical studies that mimic human responses to excessive alcohol consumption. In the current study, mice that drank 20% alcohol daily for 34-40 weeks had lower levels of muscle mass and contractile torque, indicative of alcohol-induced muscle atrophy and weakness^42,43^. Gold-standard clinical markers of ALD include elevated serum levels of AST and ALT^61^, which reflect their leakage from hepatocytes upon alcohol-induced injury^62^. While serum AST and ALT levels were unable to be quantified in the current cohort of mice due to lack of remaining sample, separate work in another cohort of C57BL/6 (same strain as present study) mice that drank 20% alcohol for ∼60 weeks found elevated serum AST and ALT levels (unpublished observations), supporting the likelihood that liver damage is already developing or progressing by the 34-40 week timepoint in our model. Moreover, in a recent large cohort study in humans, people with lower hepatic levels of GOT1 (AST) and GPT (ALT) experienced more severe ALD^63^, suggesting that declines in AST and ALT levels in liver tissue itself may be indicative of ALD development. Consistent with this, mice consuming alcohol in the present study had diminished levels of liver Got1 and Gpt (see supplemental data), as well as a lower hepatic PC:PE ratio, which is a hallmark of impaired hepatocyte membrane integrity that leads to progression of steatosis into steatohepatitis^64^ and is also indicative of ALD in both patients and animal models^45^. Thus, while we acknowledge that our current work would have benefited from liver histological analysis to confirm the true presence and severity of ALD, the observations above offer some support to the validity of our model for studying alcohol-induced dysregulation of both liver and skeletal muscle at the molecular level.

The liver metabolises over 90% of absorbed alcohol, making it highly susceptible to the toxic effects of chronic alcohol intake^12,13^. Greater molecular disruption by chronic alcohol in the liver compared to muscle is therefore logical. Consistent with this notion, we found that the liver is more sensitive than muscle to chronic alcohol at the levels of the transcriptome, proteome, and metabolome. Widespread changes across the liver transcriptome, proteome and metabolome have also been observed in human ALD models^63,65^, though omics work in human alcohol myopathic muscle tissue remains lacking. Conversely, the lipidomes of liver and muscle were found to be equally susceptible to chronic alcohol consumption. It is well established that alcohol causes gross dysregulation of liver lipids^45^. Moreover, previous human lipidomics research noted that wholesale hepatic lipidome changes associate with alcohol-related liver injury^66^. We therefore postulate that liver and muscle lipidomes exhibit similar sensitivity to chronic alcohol due to a relative increase in muscle lipidome sensitivity rather than a decline in liver lipidome sensitivity. This could be due to several factors. First, it is plausible that remodelling of the muscle lipidome after chronic alcohol intake resulted from lipid “spillover” from other sources (e.g., adipose tissue) that were then transported into muscle. Second, it is possible that alcohol directly modified muscle lipids. Indeed, many upregulated lipids in muscle (and liver) were phosphatidylethanols, which only form in the presence of ethanol^50^. Lastly, the lipidome provides a snapshot of processes across the genome, transcriptome, and proteome^67^, so cumulative changes across those layers could predispose muscle to gross lipidome remodelling. Taken together, such data indicate that lipid composition in skeletal muscle is highly influenced by chronic alcohol consumption.

Another interesting observation made in the present study was that there were few individual protein abundance changes in muscle after chronic alcohol intake, even though a substantial number of genes were found to be dysregulated. This lack of individual protein changes relative to gene changes may be due to a ‘bulk’ impairment in translational efficiency, similar to what occurs in other long-term models of muscle decline such as ageing^68^. Another plausible explanation is that muscle exhibits a ‘biphasic’ adaptive temporal response to chronic alcohol intake akin to other chronic stimuli like exercise training^69^, such that the molecular signatures captured at our end time point reflect the onset of ‘later’ molecular mechanisms of alcohol-related myopathy which are initially prominent at the transcriptome level. Alternatively, it may be that mRNA processing undergoes ‘fine-tuning’ in alcohol myopathic muscle once protein levels converge on a ‘set point’ as rates of muscle decline subdue over time^70^. In this scenario, larger changes in key mRNA could gradually couple to subtle protein-level changes that still contribute to alcohol-related myopathy. Supporting this notion, we observed a strong concordance between the transcriptome and proteome in *whole-pathway* changes related to muscle maintenance/function after chronic alcohol intake (Figure 5). Thus, heavy alcohol drinking is characterised by a more ‘global’ and persistent ‘steady-state’ mRNA-protein response in the liver than in skeletal muscle. This observation may reflect temporal differences in the onset and progression of ALD *versus* alcohol-related myopathy which might, in turn, have implications for developing optimal therapeutics for each tissue.

Arguably the most pertinent finding in the current study was that the molecular profiles induced in the liver and muscle by chronic alcohol are mostly unique to one another, implying that largely distinct mechanisms characterise ALD and alcohol-related myopathy. The molecular profile of the liver was mainly defined by an extensive and multi-layered metabolic remodelling signature, encompassing changes related to lipid, carbohydrate and glucose/glycogen metabolism that were not observed in muscle. Extensive metabolic reprogramming has also been witnessed in human ALD liver tissue omics^65^, and our findings extend to suggest that liver and muscle experience distinct metabolic reprogramming events due to chronic alcohol drinking. Widespread metabolic dysregulation is a hallmark feature of alcohol-induced liver damage. Indeed, the majority of absorbed alcohol is oxidised in the liver to acetate, during which aberrant changes in hepatic metabolism across glycolytic, gluconeogenic and fatty acid pathways occur, exacerbating the ALD phenotype^71,72^. Congruent with liver transcriptomics in ALD patients^73^, cholesterol homeostasis was strongly downregulated in liver the current work. Moreover, we found that Cyp3a family members involved in metabolism were among the genes most significantly downregulated by alcohol in liver, marrying observations of decreased hepatic CYP3A4 levels in human omics ALD research.^73,74^

Molecular profiles related to mitochondrial translation and oxidative phosphorylation were also noted to be uniquely upregulated by alcohol in the liver herein, supporting the notion that enhanced mitochondrial respiration in the liver due to alcohol consumption is associated with greater injury and damage^75^. Transcriptomics data in ALD patients also points to elevated oxidative phosphorylation gene expression in peripheral blood.^76^ Elevated hepatic mitochondrial respiration with chronic alcohol may reflect an adaptive response to enhance liver alcohol metabolism or could signify chronically ‘overworked’ mitochondria. The latter would lead to an increased mitochondrial reactive oxygen species (ROS) production^77^, which is implicated as a key mechanism in ALD^78^. Consistent with this, Glutathione S-transferase (GST) enzymes (Gstm1, Gstm3, Gstp1) were among the top uniquely upregulated molecules in the liver of mice that drank alcohol. GSTs help protect against cellular damage by detoxifying ROS and other harmful products^79^. Elevated hepatic GSTP1 abundance has recently been uncovered as a strong candidate biomarker of ALD progression in patients,^63^ with meta-analyses also demonstrating that people with *GSTM1* and *GSTP1* allelic variations hold increased susceptibility to ALD^80^. Interestingly, we also noticed Fcn1 among the top upregulated proteins by chronic alcohol specifically in liver, consistent with recent hepatic proteomic observations in patients with severe ALD and possibly linked to inflammation^81^. Extending previous studies on liver damage^82,83^, our data also revealed MTA3 as a promising new metabolism-related therapeutic target of ALD for future mechanistic investigation.

Compared to ALD, mechanistic investigations of alcohol-related myopathy are limited and mainly restricted to conventional targets of muscle protein synthesis (MPS) and breakdown (MPB)^31^. Here, in contrast to the liver, skeletal muscle was characterised by a molecular profile consistent with impaired mitochondrial energetics. After chronic alcohol consumption, muscle also displayed a unique molecular pattern primarily reflective of elevated inflammation and cytoskeletal/ECM remodelling. Chronic inflammation and mitochondrial dysfunction can both be deemed central tenets in the aetiology of muscle atrophy and weakness, potentially contributing to alcohol-related myopathy by hindering MPS pathways and/or aggravating MPB pathways^84,85^. Cytoskeletal/ECM alterations are also a major component of skeletal muscle remodelling^86^. An upregulated matrisome profile in muscle after chronic alcohol intake has been suggested to reflect a profibrotic phenotype^28^. Conversely, many matrisome features upregulated by alcohol in the present work may, in fact, be upregulated to maintain or improve muscle mass and function^87,88^. Thus, whether these muscle matrisome changes truly represent a profibrotic phenotype, or instead reflect a compensatory mechanism to avoid advanced muscle wasting with prolonged alcohol use, warrants further determination. Nonetheless, our analysis pinpoints LYL1 (an inflammatory regulator), PPARGC1a (a regulator of mitochondrial biogenesis) and CRELD1 (a matrisome member) transcription factors as key mechanistic targets of alcohol-related myopathy.

It is also intriguing to rationale how the above findings parallel the gross lipidome remodelling observed in skeletal muscle after chronic alcohol intake. On one hand, inflammation and mitochondrial dysregulation are prime events that can influence muscle lipid dynamics^89,90^. Yet, the lipidome alterations we found appear underscored by wholesale changes in phospholipids, particularly PC and PE species. Muscle phospholipids have been linked to declines in muscle size and strength^91,92^, albeit via unknown mechanisms, and PC:PE dynamics are implicated in promoting mitochondrial dysfunction and inflammation^44,93^. Phospholipids are also integral components of the sarcolemma^94^, suggesting that muscle PC:PE ratio may be inherently linked to cytoskeletal/ECM remodelling. Thus, our current findings support our recent postulation that remodelling of muscle phospholipid composition may play a role in the aetiology of alcohol-related myopathy^95^ – potentially through some form of mitochondria-inflammatory-matrisome regulatory circuit. Indeed, network modelling uncovered two multi-omic sub-networks in muscle after alcohol intake, both of which had a central ‘hub’ phospholipid connected to cytoskeletal/ECM, inflammation and/or mitochondria-related transcripts. Coined the ‘lipotranscriptome’, lipid regulation of the transcriptome is emerging as a promising target for the discovery and development of disease diagnostics and therapeutics^96^. Alcohol-induced changes in muscle lipotranscriptome that centre on phospholipid regulation of mitochondria, inflammation and/or the matrisome may, therefore, be among the strongest candidates for understanding and treating chronic alcohol-related myopathy.

Traditional drug discovery relying on lab-based compound screens is cumbersome, costly, labour-intensive and high-risk^97^. Omics circumvents many of these barriers to accelerate the drug discovery process by enabling *in silico* drug repurposing on an unprecedented scale^98^. Exploiting this approach, we used human drug signature databases to computationally predict a refined list of compounds that target molecular profiles of chronic alcohol-related liver and/or muscle. While the interpretation of all predicted compounds is beyond the scope of this discussion, this list provides a useful tool to expedite future hypothesis-driven work on ALD and alcohol-related myopathy therapeutics. For example, among the most prevalent compounds predicted to target the unique molecular profile of chronic alcohol-related liver were saracatinib and GSK126. Saracatinib has been reported to attenuate liver fibrosis by preventing activation of hepatic satellite cell activation^56,99^, while GSK126 also appears to have anti-fibrotic liver properties^55^ and decreases liver fatty acid content^100^. These two compounds may thus offer promising avenues for exploration in the context of treating ALD. We also identified metformin and trichostatin A as strong candidate therapeutics for alcohol-related myopathy. Indeed, trichostatin A has been shown to ameliorate muscle atrophy induced by unloading^58^ and to reduce muscle fatty acid infiltration^101^. While the muscle therapeutic potential of metformin is somewhat equivocal, evidence indicates that it could help counter muscle decline in pathological scenarios, as would be excessive chronic alcohol drinking, potentially by normalising mitochondrial dysfunction or disturbed energy metabolism^59^. Further, compounds that target the molecular profiles of both alcohol-related muscle and liver might represent the most viable concurrent treatments for ALD and alcohol-related myopathy. Notably, MERCK544 may be a viable compound for dual-therapeutic targeting of ALD and alcohol-related myopathy by mitigating 11β-HSD1-dependent metabolic dysregulation related to both liver and muscle^102–104^.

In summary, we undertook the first multi-omic screening of liver *versus* skeletal muscle responses to chronic alcohol use. Our results provide several new insights into therapeutic targets of and pharmacologic interventions for ALD and alcohol-related myopathy. Nonetheless, comprehensive follow-up investigations remain crucial to validate these mechanistic lines of inquiry and to assess the true efficacy of our predicted drug compounds for mitigating alcohol-induced liver and muscle maladaptations. Such work should include tracking the time course of multi-omic responses to chronic alcohol in both sexes and incorporate a wider range of physiological and morphological readouts (e.g., liver histological analysis to assess stage of liver injury), to provide a more complete picture of the mechanisms underpinning the onset and progression of ALD and alcohol-related myopathy. Thorough examination of predicted drug compounds, including optimal dosing, delivery strategies and safety is also warranted, at both the individual tissue and cross-tissue levels. Overall, our current findings provide a strong benchmark for expediting the mechanistic understanding of ALD and alcohol-related myopathy in humans and may help accelerate the development of optimal personalised countermeasures.

## Materials and methods

### Experimental overview

The mice and experimental procedures used are as previously reported^42,43^. Briefly, female C57BL/6 mice obtained from Jackson Laboratory were aged to 23-28 weeks and then randomly assigned access to either a 100% water bottle (Control mice; *n* = 9) or a bottle containing 80% water + 20% alcohol (ethanol) (Alcohol mice; *n* = 14) for 34-40 weeks. Female mice were used as it has been reported that females are more susceptible to alcohol-related myopathy and ALD^105–107^. Alcohol-consuming mice were acclimatised to alcohol by increasing alcohol concentration in 5% increments from 0% to 20% (w/v) across two weeks. All mice were supplied standard rodent chow *ad libitum* throughout. At study completion, body composition was assessed using the Bruker Minispec NMR Analyzer (LF50 Series, model mq 7.5), *in vivo* plantarflexor muscle isometric torque using a servomotor system (Model 300C-LR and 701C; Aurora Scientific, Aurora, Ontario, Canada), and blood alcohol concentration using an AM1 alcohol analyser (Analox Instruments Ltd.). Mice were euthanised under anaesthesia (2-to-3% isoflurane) by exsanguination followed by cervical dislocation, then plantarflexor muscles and liver harvested, weighed, snap frozen in liquid nitrogen and stored at -80°C until further analyses. For the purposes of this work, a composite of blood alcohol concentration, body composition, and muscle mass and torque data from two previous publications has been included^42,43^. Experimental procedures were approved by the Ohio University Animal Care and Use Committee. All methods were performed in accordance with the relevant guidelines and regulations. The study is reported in accordance with ARRIVE guidelines.

### Omics data generation

Harvested liver and muscle was shipped on dry ice to BGI Americas (San Jose), where tissue samples were processed, and omics data generated. For transcriptomics, strand-specific (second strand cDNA synthesis with dUTP) 100 bp paired-end reads were generated from extracted RNA using the DNBseq platform. For proteomics, a label-free quantitative approach was undertaken using nano flow HPLC (Ultimate 3000) followed by Orbitrap Eclipse Tribrid Mass Spectrometer (Thermo Fisher Scientific, USA). A spectral library was constructed via data-dependent acquisition using the MS2-based method, using a fractionated composite of all digested samples. Data-independent acquisition was subsequently performed on each sample via a high-resolution full mass spectrometry scan followed by two data-independent acquisition segments. Raw mass spectrometer files were input into MaxQuant (v1.5.3.30; https://www.maxquant.org/) for identification and quantification against the mouse database, with identified peptides that satisfied a false discovery rate ≤ 1% used when constructing the final spectral library. For metabolomics and lipidomics, features were extracted from tissue samples using solvent-based precipitation, with feature separation and detection performed using a UPLC I-Class Plus (Waters, USA) tandom Q Exactive high resolution mass spectrometer (Thermo Fisher Scientific, USA), utilising AQUITY UPLC BEH C18 and Amide columns for metabolites and a CSH C18 column for lipids (Waters, USA). In any case, QC samples were produced by pooling equal volumes of prepared supernatant (10μL) from each sample. Off-line mass spectrometry data were subsequently input into Compound Discover (v3.3; Thermo Fisher Scientific, USA) for metabolite peak extraction and identification (using BGI metabolome, mzcloud and chemspider databases as reference) or LipidSearch (v4.1; Thermo Fisher Scientific, USA) for lipid peak extraction and identification.

### Omics data pre-processing

For transcriptomic data, reads were cleaned using SOAPnuke software (v2.2.1; https://github.com/BGI-flexlab/SOAPnuke) to filter out adaptor sequences, contamination and low-quality reads^108^. The clean reads were then aligned to the mouse reference transcriptome using Kallisto (v0.46.1, with bias correction; https://pachterlab.github.io/kallisto/)^109^. The reference transcriptome was compiled from the Ensembl Release 109 mouse reference genome (primary assembly) and associated transcript annotations using GFFread (v0.12.7; https://github.com/gpertea/gffread)^110^. Gene counts were inferred from transcript abundance estimates scaled to library size using the tximport R package (v1.28.0; https://doi.org/doi:10.18129/B9.bioc.tximport)111. Lowly expressed genes were then removed using the limma R package (v3.56.2; https://doi.org/doi:10.18129/B9.bioc.limma) *filterByExpr* function (with tissue as the grouping factor and ‘min.prop’ set to 1), and counts for remaining genes (*n* = 14,017) normalised using the limma-voom approach as per developer guidelines for mixed-design studies. For proteomic data, log2 transformation was applied, proteins with more than 50% missing values omitted and the k-Nearest Neighbour Algorithm used to impute missing values, resulting in normalised abundances for *n* = 3,512 proteins. For metabolomics data, each UPLC column’s Compound Discover output was subject to probabilistic quotient normalisation to obtain relative peak areas, quality control-based robust LOESS signal correction to correct batch effect, removal of features with a coefficient of variation larger than 30% based on relative peak area in QC samples, filtering for features with a recognised KEGG ID, and log2 transformation. Normalised metabolite abundances for each column were then aggregated based on mean value to produce a singular metabolomic dataset (*n* = 873 features). For lipidomic data, the LipidSearch output was subject to removal of lipids with > 50% missing values in QC samples and > 80% missing values in experimental samples, imputation of remaining empty values via the k-Nearest Neighbour Algorithm, probabilistic quotient normalisation to obtain relative peak areas, quality control-based robust LOESS signal correction to correct batch effect, filtering to remove features with a coefficient of variation larger than 30% based on relative peak area in QC samples, and log2 transformation, leading to normalised abundances of *n* = 744 lipids. Prior to log2 transformation, percent content of PC and PE lipid classes in each sample (sum of relative peak values for a given class in the sample, divided by total of all relative peak values in the sample) were also calculated and used to quantify muscle and liver PC:PE ratios.

## Statistics

### Statistical evaluation of end-point data

End-point variables (total body mass, lean mass, fat mass, plantarflexor muscle mass, plantarflexor muscle torque, liver mass, liver-to-body mass ratio, plantarflexor muscle PC:PE ratio, liver PC:PE ratio, blood alcohol concentrations) were compared between alcohol-consuming mice and control mice on a per-tissue basis using either the two-tailed Student’s independent t-test (when normal distribution and equal variance), the two-tailed Welch’s independent t-test (when normal distribution but unequal variance) or the Mann-Whitney U test (when non-normal distribution). Analyses were conducted in R (v4.3.1; https://www.r-project.org/), with statistical significance accepted when P ≤ 0.05. Unless otherwise stated, in-text descriptive statistics are presented as median (inter quartile range).

### Omics differential expression analysis

Differential analysis was undertaken at the level of each individual omic strand using the limma R package (v3.56.2; https://doi.org/doi:10.18129/B9.bioc.limma)112. Briefly, linear mixed effects models were fitted with condition (Alcohol, Control) as a ‘fixed’ effect and mouse ID as a ‘random’ effect, utilising the developers’ recommended duplicateCorrelation approach. The empirical Bayes method was then used to calculate moderated t-scores^113^, and comparisons between Alcohol and Control mice per tissue extracted. For each omics layer, features with a Benjamini-Hochberg adjusted P-value ≤ 0.1 were defined as being differentially regulated by chronic alcohol. Differentially regulated features were then overlaid to identify features commonly or uniquely dysregulated by chronic alcohol in liver versus muscle.

Omics over-representation analysis of pathways, TF targets and molecular classes: Features commonly or uniquely regulated by chronic alcohol in the liver versus muscle were subject to over-representation analyses of pathways, TF targets and molecular classes using the clusterProfiler R package (v4.8.3; https://doi.org/doi:10.18129/B9.bioc.clusterProfiler) enrichr function^114^. For transcripts and proteins, analyses were performed against MSigDB (v2023.1)^115^ mouse Molecular Hallmark, Reactome Pathway and TF target (GTRD) gene sets. Default values for minGSSize and maxGSSize arguments were used, except for GTRD sets, where maxGSSize was set unbounded. For metabolites, analyses were performed against final class, super pathway and sub pathway sets as assigned during metabolite identification (minGSSize = 3, maxGSSize unbounded). For lipids, analysis was performed against lipid main class sets as assigned during lipid identification (minGSSize = 3, maxGSSize unbounded). For each omic strand, the corresponding background list contained all annotated features utilised during differential testing. Over-represented sets were selected as those with a Benjamini-Hochberg corrected P-value ≤ 0.05 that were enriched for at least 2 given features.

### Rank-based analysis across the transcriptome and proteome

The rank-rank hypergeometric overlap method^48^ was employed (via the RRHO2 R package, v1.0; https://github.com/RRHO2/RRHO2) to decipher the general degree of correspondence between differential gene expression and protein abundance patterns in the liver and in muscle. The algorithm was applied to unique features present at both the gene level and protein level (n = 3,303 genes/proteins), with features ranked by t-score. Gene set enrichment analysis was also employed as a rank-based method to elucidate global pathway regulation at the gene level and protein level in the liver and in muscle. These analyses were performed using the clusterProfiler R package (v4.8.3; https://doi.org/doi:10.18129/B9.bioc.clusterProfiler) GSEA function against MSigSB Molecular Hallmark and Reactome Pathway gene sets as above (default minGSSize and maxGSSize argument values). In each case, the algorithm was applied to unique features present at both the gene level and protein level (n = 3,303 genes/proteins), with features ranked by t-score and enrichment defined at the Benjamini-Hochberg corrected P-value ≤ 0.1 level.

### Multi-omic relevance network analysis

Features from all omics layers were integrated using multi-omic correlation network analysis with the Mixomics R package (v6.24.0; https://doi.org/doi:10.18129/B9.bioc.mixOmics) DIABLO method^52^. Multiblock integration with projection to latent structure models with discriminant analysis was performed, treating each omic strand as a block and condition as the discriminator. A design matrix was used to maximise the strength of relationships between blocks. Relevance networks were extracted for strongly correlated features across omics layers (|correlation coefficient| > 0.85). Component 1 was chosen for association estimates in the liver and component 2 for muscle, based on inspection of individual sample plots and loading weights. Sub-networks of each relevance network were determined via multi-level community analysis using the igraph R package (v1.5.1; https://r.igraph.org/)^116^, with |correlation coefficient| used as edge weight. Highly connected ‘hub’ features of each sub-network were defined with an eigenvector centrality score > 0.7. Networks were visualised using Cytoscape (v3.10.0; https://cytoscape.org/)^117^.

### Omics-driven drug prediction analysis

Transcriptome and proteome features commonly or uniquely dysregulated by chronic alcohol in the liver versus muscle were further subjected to over-representation analyses against human Drug Signature Database^53^ and Proteome Drug Atlas^54^ sets using the Enrichr online webtool (https://maayanlab.cloud/Enrichr/)^118^. For each omic strand, the background list contained all annotated features utilised during differential testing. Ortholog conversions were handled implicitly as part of the built-in functionality of the Enrichr tool. Over-represented sets were selected as those with a Benjamini-Hochberg corrected P-value ≤ 0.05 that were enriched for at least 2 given features.

## Data availability

Raw transcriptomic data from current study is deposited in the Sequence Read Archive repository (https://www.ncbi.nlm.nih.gov/sra) with links to BioProject ID PRJNA1274880. Raw proteomic data from current study is deposited in the EBI Proteomics Identification Database repository (https://www.ebi.ac.uk/pride/) under the dataset identifier PXD064997. Raw metabolomic and lipidomic data are deposited in the EBI MetaboLights repository (https://www.ebi.ac.uk/metabolights/) under the study identifier MTBLS12616. Underlying values of the data reported in the manuscript itself are provided as supplementary material in Document S1.

## Author contributions

Conceptualisation: C.R.G.W. & C.W.B.; Mouse experimental procedures: M.S.G., A.M.B., S.E.M. & C.W.B.; Data analysis: C.R.G.W.; Figures and visualisation: C.R.G.W.; Manuscript original draft: C.R.G.W.; Manuscript review, editing and approval: C.R.G.W., M.S.G., A.M.B., S.E.M., N.J.S., B.C.C. & C.W.B..

## Acknowledgements

C.R.G.W. and C.W.B. acknowledge support from the UK Research and Innovation Doctoral Career Development Fund scheme, which helped to fund this work. This work was also funded, in part, through the University of Exeter’s Project ADA (Accelerating Data Science and Artificial Intelligence). N.J.S. acknowledges the support of the Osteopathic Heritage Foundation through funding for the Osteopathic Heritage Foundation Ralph S. Licklider, D.O., Research Endowment in the Heritage College of Osteopathic Medicine. B.C.C. acknowledges the support of the Osteopathic Heritage Foundation through funding for the Osteopathic Heritage Foundation Harold E. Clybourne, D.O., Endowed Research Chair in the Heritage College of Osteopathic Medicine. C.W.B. acknowledges the support of the Osteopathic Heritage Foundation through funding for the Osteopathic Heritage Foundation Ralph S. Licklider, D.O., Endowed Faculty Fellowship in the Heritage College of Osteopathic Medicine. This work was also supported, in part, by grants from the National Institutes of Health (R01AG044424 & R01AG067758 to B.C.C.). The authors thank Thomas Krauss for assisting with revisions.

## Supplementary Information Files

**Figure S1:** Reactome Pathways enriched with genes dysregulated by chronic alcohol in the liver and/or muscle. Bubble plot depicts results from over-representation analysis of MSigDB mouse Reactome Pathway gene sets for genes commonly/uniquely dysregulated by chronic alcohol use in the liver and muscle. Circle size is proportional to the number of gene hits as a % of the total number of annotated genes for a given overlap. Red and blue shading denote significant over-representation in upregulated and downregulated genes, respectively (adjusted *P* ≤ 0.05, enriched for ≥ 2 genes).

**Document S1:** A Single XLSX file containing underlying values of reported data.

