## Supplementary figures and images for "Chronic alcohol intake elicits distinct multi-omic profiles in the liver *versus* skeletal muscle of mice"

### Supplemental Figure 1

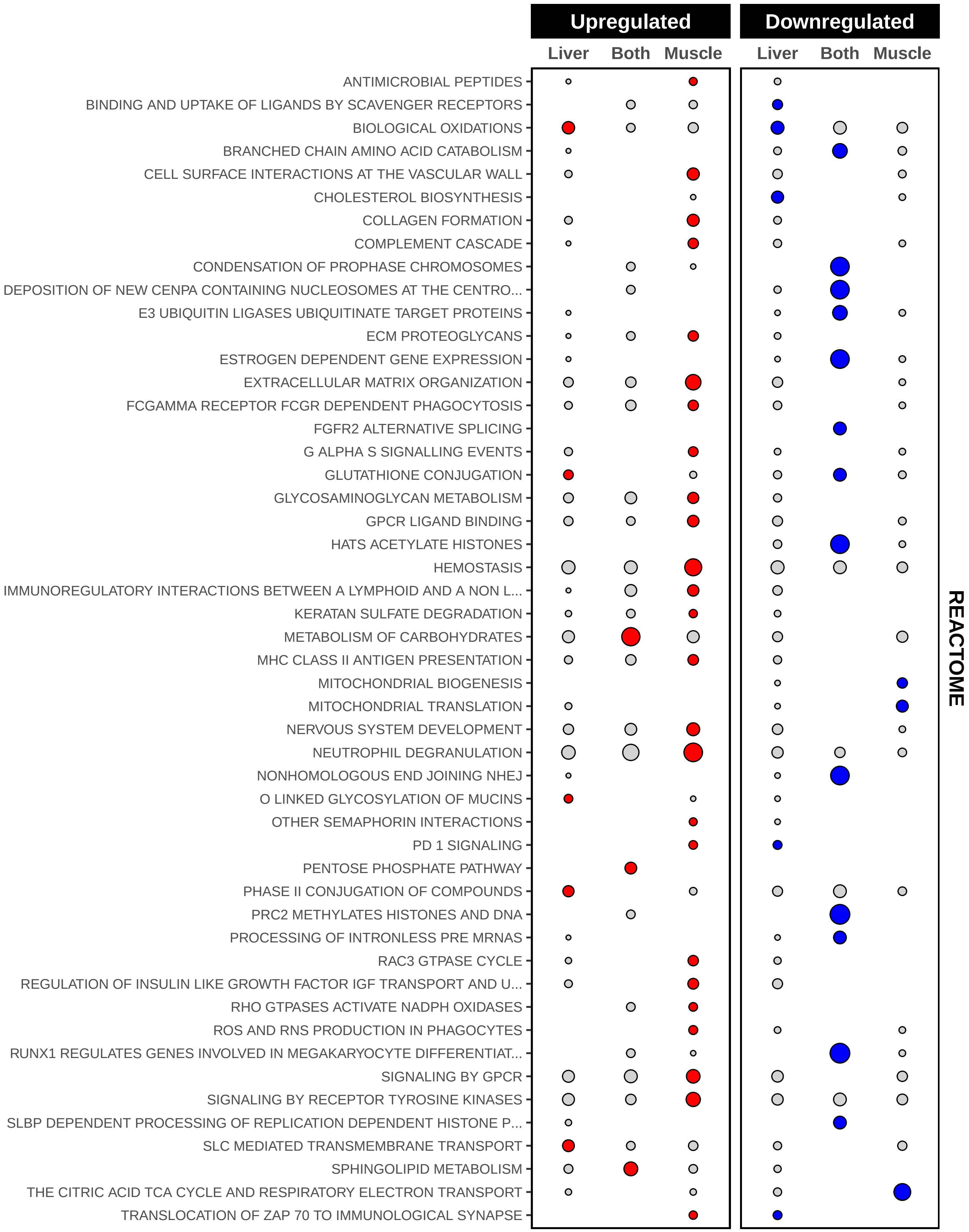
